# The jasmine (*Jasminum sambac*) genome and flower fragrances

**DOI:** 10.1101/2020.12.17.420646

**Authors:** Gang Chen, Salma Mostafa, Zhaogeng Lu, Ran Du, Jiawen Cui, Yun Wang, Qinggang Liao, Jinkai Lu, Xinyu Mao, Bang Chang, Li Wang, Zhichao Jia, Xiulian Yang, Yingfang Zhu, Jianbin Yan, Biao Jin

**Affiliations:** College of Horticulture and Plant Protection, Yangzhou University, Yangzhou 225009, China; College of Bioscience and Biotechnology, Yangzhou University, Yangzhou 225009, China; Shenzhen Branch, Guangdong Laboratory for Lingnan Modern Agriculture, Genome Analysis Laboratory of the Ministry of Agriculture, Agricultural Genomics Institute at Shenzhen, Chinese Academy of Agricultural Sciences, Shenzhen 518120, China; College of Landscape Architecture, Nanjing Forestry University, Nanjing 210037, China; Institute of Plant Stree Biology, State Key Laboratory of Cotton Biology, Department of Biology, Henan University, Kaifeng 475001, China

## Abstract

*Jasminum sambac*, a world-renowned plant appreciated for its exceptional flower fragrance, is of cultural and economic importance. However, the genetic basis of its fragrance is largely unknown. Here, we present the first *de novo* genome of *J. sambac* with 550.12 Mb (scaffold N50 = 40.1 Mb) assembled into 13 pseudochromosomes. Terpene synthase genes associated with flower fragrance are significantly amplified in the form of gene clusters through tandem duplications in the genome. Eleven homolog genes within the SABATH super-family were identified as related to phenylpropanoid/benzenoid compounds. Several key genes regulating jasmonate biosynthesis were duplicated causing increased copy numbers. Furthermore, multi-omics analyses identified various aromatic compounds and the key genes involved in fragrance biosynthesis pathways. Our genome of *J. sambac* offers a basic genetic resource for studying floral scent biosynthesis and provides an essential foundation for functional genomic research and variety improvements in *Jasminum*.

## Introduction

*Jasminum sambac* (common names: Arabian jasmine, Sambac jasmine, jasmine flower, 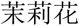 Mo-Li-Hua) is famous worldwide as a fragrant plant with sweet-scented flowers. The fragrant flowers of *J. sambac* are used for the extraction of jasmine essential oil, which is a common natural ingredient in the perfume and cosmetic industries, as well as in pharmaceutical applications and aromatherapy^1,2^. *J. sambac* flowers are also used in the manufacture of jasmine tea consumed popularly in East Asia^^3^,4^. Various food products with the sweet flavor of *J. sambac* flowers have been produced, such as syrup, aerated water, jam, yogurt, ice cream, wine, etc. In some Asian countries, *J. sambac* is regarded as auspicious symbols in religious ceremonies or used to make garlands for welcoming guests^5^ and it has been integrated into local cultures and traditions^6,7^.

Flower fragrances are volatile organic compounds (VOCs) emitted by flowering plants to attract pollinators and ensure reproductive success^8^. Flower fragrances also attract humans and have become the focus of intensive use in the floriculture and fragrance industries. Different flowering plant species have distinct flower fragrances, depending on differences in the composition, amount, and emission of floral VOCs^8,9^. The VOCs of *J. sambac* floral scents belong mainly to the terpenoid and phenylpropanoid/benzenoid classes^10^. However, most previous analyses of VOCs from *J. sambac* flowers were based on harvested flowers^11–13^, whereas the fragrances actively released by flowers growing in a natural state remain obscure. Some genes involved in the biosynthetic pathways of *J. sambac* floral scent compounds have been isolated and analyzed, such as genes responsible for the biosynthesis of α-farnesene (*JsHMGS*, *JsHMGR*, *JsFPPS*, and *JsTPS*) in the mevalonic acid (MVA) pathway^14^. However, the biosynthesis pathways of floral scent compounds and their regulatory networks are complex and their underlying genetic mechanisms remain largely unknown. Whole-genome sequencing is a practical strategy for identifying the metabolic pathways of natural-compound biosynthesis in plants^15–17^. Although *J. sambac* flower products are widely used and its flower scents are economically valuable, the lack of *J. sambac* genome data seriously hampers progress in unraveling its fragrance biosynthesis and metabolism. Additionally, jasmonates are important aromatic substances in *Jasminum* flowers. Jasmonates have been extensively studied in biotic and abiotic stress responses and defenses in model plants, crops, and other plants^18^. Nevertheless, research on jasmonate biosynthesis and regulation in *Jasminum* is also impeded by the absence of genome sequence data.

Here, we report a chromosome-level genome assembly of *J. sambac* obtained using a combination of Illumina and PacBio data, enhanced by information from Hi-C technologies. Furthermore, by combining multi-omics analyses of different stages of flowers, we identified various aromatic compounds released from both harvested and naturally grown flowers. Several important genes involved in the biosynthetic pathways of major fragrant compounds in jasmine flowers were identified. This *J. sambac* genome sequence and the identified floral scent volatiles offer valuable resources for *J. sambac* genetic research and will lay a foundation for biological and agronomic research on this commercially and culturally important species.

## Results

### Genome sequencing and assembly

K-mer analysis of parallel next-generation sequencing short-read data revealed that the *J. sambac* genome size is ~573.02 Mb with a heterozygosity rate of 0.99% and a repeat rate of 57.12% (Supplementary Fig. S1), indicating the complexity of the *J. sambac* genome. The genome was sequenced and assembled using a combination of single-molecule real-time (SMRT) sequencing technology from PacBio and Hi-C. In total, 63.82 Gb of data (116× the assembled genome) were generated from 4.5 M PacBio single-molecule long reads (average read length = 14.2 kb, longest read length = 127.0 kb). In addition, using DNA from the leaves of *J. sambac*, we generated 61.6 Gb of Illumina paired-end reads (~112×). PacBio long reads were assembled using the overlap-layout-consensus method with different assemblers. Finally, *de novo* assembly yielded 373 contigs with a contig N50 length of 2.50 Mb. The total assembly size was 550.12 Mb, with a GC content of 34.62%, covering 96% of the estimated *J. sambac* genome size (Table 1).

**Table 1.** Statistics of the *J. sambac* genome and gene annotation

| Genome assembly |  |
| --- | --- |
| Estimated genome size | 573.02 Mb |
| GC content | 34.62% |
| Contig N50 length | 2.50 Mb |
| Longest contig | 7.64 Mb |
| Assembled genome size | 550.12 Mb |
| Scaffold N50 length | 40.10 Mb |
| Longest scaffold | 52.97 Mb |
| Gene annotation |  |
| Repeat region | 47.22% |
| Number of protein coding genes | 30,129 |
| Average transcript length | 3335.86 bp |
| Average coding sequence length | 1103.73 bp |
| Average exons per gene | 4.87 |
| Average exon length | 226.57 bp |
| Average intron length | 576.58 bp |

To refine the *J. sambac* assembly, Hi-C libraries were constructed and sequenced. The Hi-C read pairs were mapped onto the draft assembly and used to improve the scaffold N50 to 40.1 Mb and the contig number to 383, with the longest scaffold being 53.0 Mb and the scaffold number being 112 (Table 1, Supplementary Table S1). The final reference assembly comprised 13 chromosome-scale pseudomolecules (the pseudomolecules are hereafter referred to as chromosomes) (Fig. 1), with maximum and minimum lengths of 53.0 Mb and 33.5 Mb, respectively (Supplementary Table S2). The total length of the chromosomes accounts for 97.36% (535.57 Mb) of the assembled genome size of 550.12 M.

**Fig. 1.**
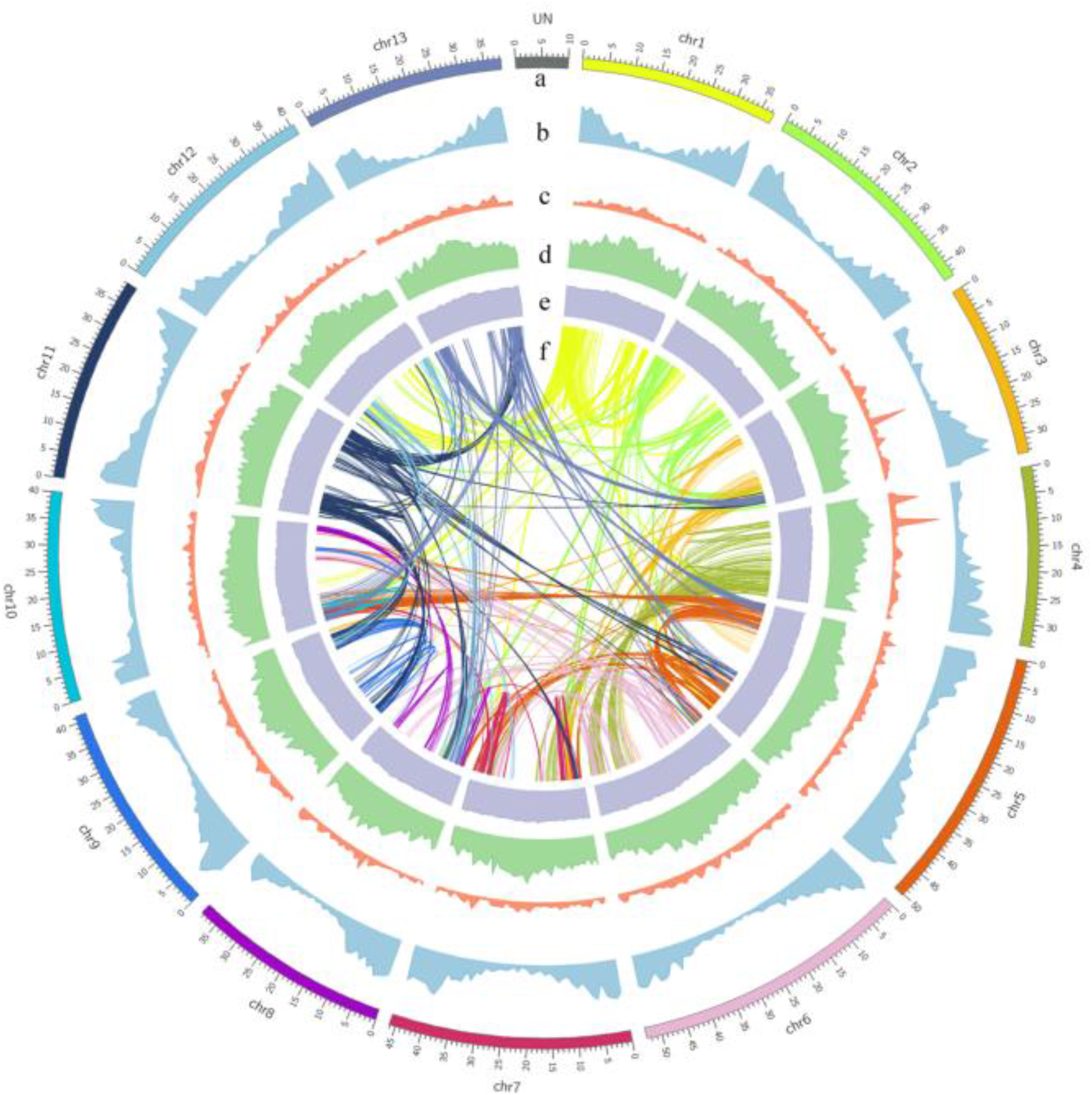
Genomic features of *J. sambac*. **a** Circular representation of the 13 pseudochromosomes. **b** Gene density. **c** Density of non-coding RNA. **d** Distribution of transposable elements (TEs). **e** GC content distribution. **f** Syntenic relationships among duplication blocks containing more than 13 paralogous gene pairs.

To evaluate the quality of the assembled genome, the Benchmarking Universal Single-Copy Orthologs (BUSCO)^19^ assessment was conducted and the results revealed a high-quality draft genome covering 1320 (91.7%, Supplementary Table S3) complete single-copy orthologs of the 1440 plant-specific sequences (Embryophyta data set from BUSCO datasets). Furthermore, a Hi-C interaction heatmap separated distinct regions on different chromosomes, indicating that all bins were allocated to 13 chromosomes (Supplementary Fig. S2). These results demonstrated that the assembled *J. sambac* genome is of high quality at the chromosome level.

### Genome annotation and gene prediction

We identified a total of 259.8 Mb of repetitive sequences in the genome of *J. sambac*, which accounted for 47.22% of the assembled genome. Among them, transposable elements (TEs) were the predominant components (45.56% of the genome) and long terminal repeat (LTR) retrotransposons comprised 33.97% of the assembled genome (Supplementary Table S4). Within the LTR family, the *copia* subfamily was the most abundant, accounting for 16.2% of the genome, followed by the *gypsy* subfamily (15.0%). Additionally, the distribution of TEs varied across the genome (Fig. 1d, Supplementary Fig. S3); for example, the TE content was higher near the centromeres compared to other parts of the chromosomes.

We annotated the remaining repeat-masked *J. sambac* genome using a comprehensive strategy of *de novo* prediction combined with homology-based and transcriptome-based protein predictions. In total, 30,129 complete genes were predicted, with an average transcript length of 3336 bp and an average coding sequence length of 1104 bp (Table 1, Supplementary Tables S4, S5, S6). Among the predicted genes, 67.5% (20,345 of 30,129) were predicted by all three strategies (Supplementary Fig. S5). Additionally, most genes were distributed near the two ends of the chromosomal arms (Fig. 1b).

In total, 9902 non-coding RNAs, including 1657 microRNAs (miRNAs), 1767 ribosomal RNAs (rRNAs), and 535 transfer RNAs (tRNAs), were identified (Supplementary Table S7). Further functional annotation revealed that 93.20% of all predicted genes could be annotated with the following protein-related databases: RefSeq non-redundant database (NR) (92.80%), Swiss-Prot (74.20%), Kyoto Encyclopedia of Genes and Genomes (KEGG) (69.60%), InterPro (78.00%), Gene Ontology (GO) (53.80%), and Pfam (73.00%). In total, 18,911 genes were commonly annotated in the Swiss-Prot, InterPro, NR, and KEGG databases (Supplementary Table S8, Supplementary Fig. S6).

### Genome evolution of *J. sambac*

The evolutionary dynamics of gene families were analyzed by comparing the *J. sambac* genome with those of 16 representative plant species. In total, 42,577 gene families were clustered among all 17 species, and 6337 gene families were in common, including 3670 single-copy orthologs (Fig. 2d). From the gene families clustered in five species of the Oleaceae family (*J. sambac*, *Osmanthus fragrans*, *Fraxinus excelsior*, *Olea europaea*, and *Olea oleaster*), 15,160 gene families were identified in the *J. sambac* genome, of which 800 gene families (2060 genes) were *J. sambac*-specific while 12,037 gene families were shared among all five species in the family (Fig. 2a). Functional enrichment analysis of the *J. sambac*-specific gene families indicated that these gene families are mainly involved in terpenoid backbone biosynthesis, monoterpenoid biosynthesis, and protein processing in the endoplasmic reticulum (Supplementary Fig. S7), which are likely important for volatile compound biosynthesis in *J. sambac* flowers^10^. A phylogenetic tree was constructed from single-copy gene families of *J. sambac* and the 16 representative plant species (Fig. 2c). The results revealed that the Oleaceae and Labiatae split ~78.5 million years ago (Mya), whereas *J. sambac* diverged ~48.8 Mya from the common ancestor of the five species within the Oleaceae family. Among the Oleaceae, *J. sambac* diverged earlier than the other four species. In addition, we found 16 expanded gene families and 8 contracted gene families in *J. sambac* compared to the common ancestor of Oleaceae and Labiatae (Fig. 2b). The expanded gene families were involved mainly in riboflavin metabolism and butanoate metabolism (both related to fragrant volatiles). One contracted gene family was related to plant–pathogen interactions. We applied a four-fold synonymous third-codon transversion (4DTv) estimation to detect whole-genome duplication (WGD) events. The results revealed that one WGD event might have occurred in the common ancestor of *J. sambac* and *O. fragrans* before their divergence (Fig. 2e).

**Fig. 2.**
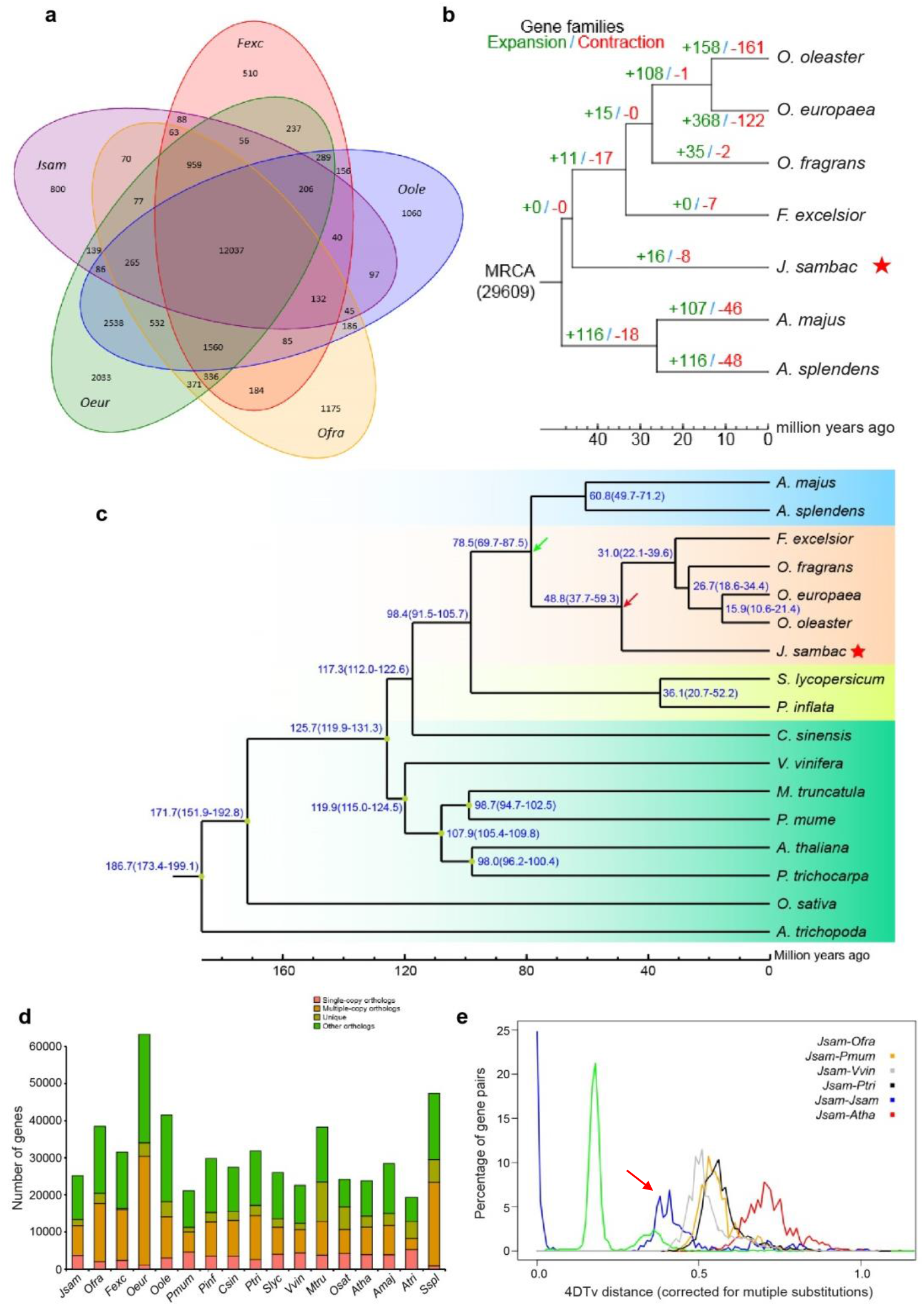
Comparative genomic analysis of *J. sambac* and other species. **a** Venn diagram of the shared orthologous gene families in *J. sambac*, *F. excelsior*, *O. fragrans*, *O. europaea*, and *O. oleaster.* The number of gene families is listed for each component. **b** Expansion and contraction in gene families. The numeric value beside each node shows the number of expanded (green) and contracted (red) gene families. **c** Phylogenetic tree constructed from single-copy gene families of *J. sambac* and 16 representative plant species. The blue numbers beside each node indicate the divergence time of each species. **d** Distribution of genes in *J. sambac* and 16 representative plant species. Only the longest isoform for each gene was used. Gene clusters (families) were identified using the OrthoMCL package with default parameters. **e** Distribution of 4DTv distance between syntenic orthologous genes. The abscissa represents the 4DTv value; the ordinate represents the proportion of genes corresponding to the 4DTv values. The red arrow indicates a WGD event that occurred before the divergence of *J. sambac* and *O. fragrans*.

### The terpene synthase (TPS) gene family and terpene biosynthesis in *J. sambac*

The TPS family is a vital enzyme gene family for terpene biosynthesis, which is crucial in the production of floral VOCs. We identified 59 TPS genes in *J. sambac* containing at least one conserved domain, and most of the TPS genes (47 of 59) contained two conserved domains (Supplementary Table S9). We further constructed an evolutionary tree of the 47 TPS genes containing two conserved domains and found that the TPS genes of *J. sambac* could be classified into five subgroups: TPS-a, TPS-b, TPS-c, TPS-e/f, and TPS-g. TPS-a was the largest subgroup, accounting for 53.2% of the total TPS genes (Fig. 3a). Furthermore, most of the TPS genes were highly expressed in leaves and flowers in *J. sambac*. The number of *J. sambac* TPS genes containing two conserved domains is significantly higher compared to *Arabidopsis thaliana* (33), *Camellia sinensis* (30), *Solanum lycopersicum* (33), *O. fragrans* (40) (Fig. 3b), cacao (36), and kiwifruit (34). Almost half of the *J. sambac* TPS genes (23 of 47) contained tandem repeats, and these genes formed TPS gene clusters on chromosomes 2, 3, 4, 6, and 11 (Fig. 3c, red gene IDs). These genes underwent recent tandem duplication events, rather than a WGD event, resulting in the amplification of TPS genes in the *J. sambac* genome (Fig. 3d). Through phylogenic analysis of the TPS genes, we further identified 17 gene pairs, 11 of which had a synonymous substitution rate (*Ks*) < 0.2 (Fig. 3e), implying that a negative selection occurred in these conservative TPS genes. Notably, we also found several events of 4:1 or 2:1 double replication of TPS genes between *J. sambac* and the tomato genome (Supplementary Fig. S8b), indicating the expansion of the TPS family of *J. sambac* (Oleaceae) relative to tomato (Solanaceae). In addition, the expression of most TPS genes was higher in flowers than in leaves (Fig. 3a), indicating that the TPS genes in *J. sambac* are functionally importantly in flowers. Furthermore, the differentially expressed genes between the full-bloom flowers (FFs) and flower buds (FBs) were enriched in several categories: terpenoid backbone biosynthesis, ubiquinone and other terpenoid-quinone biosynthesis, fatty acid metabolism, and flavonoid biosynthesis (Supplementary Fig. S8a, arrows). The mean expression of TPS genes was higher in FBs than in FFs (Fig. 3f), indicating that these genes are actively expressed at the bud stage, preparing for the release of floral fragrance substances at the full-bloom stage.

**Fig. 3.**
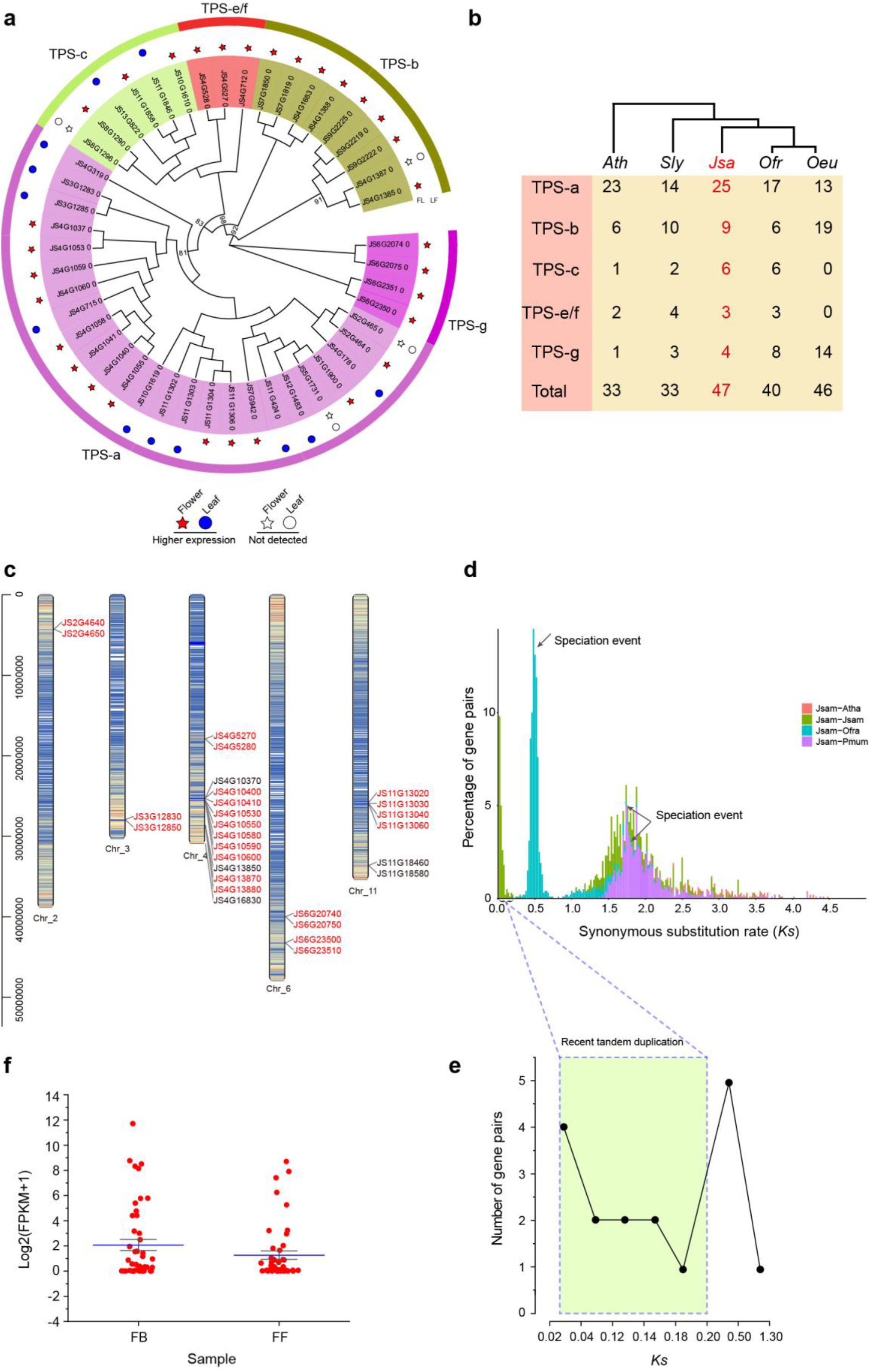
TPS gene family in the genome of *J. sambac*. **a** Evolutionary tree of the 47 TPS genes containing two conserved domains identified in the *J. sambac* genome. Colored stars and circles indicate the TPS genes highly expressed in flowers and leaves, respectively. **b** The numbers of TPS genes containing two conserved domains in *J. sambac*, *A. thaliana*, *S. lycopersicum*, and *O. fragrans*, and *O. europaea*. **c** Chromosomal distribution of the *J. sambac* TPS genes. The colored lines in different chromosomes indicate the gene density; darker lines indicate higher gene density. Red gene names indicate TPS genes that formed gene clusters on chromosomes. **d** No WGD event was identified in the amplification of TPS genes in the *J. sambac* genome. **e** *Ks* distribution of the TPS genes in the *J. sambac* genome. The TPS gene pairs in the box had *Ks* values < 0.2. **f** Expression levels of TPS genes in flower buds (FB) and full-bloom flowers (FF) of *J. sambac.* Blue bars indicate mean expression levels.

The terpene biosynthesis pathway is another important floral-fragrance pathway. We therefore examined the terpene biosynthesis pathways and confirmed that large numbers of TPS genes were involved in the synthesis of terpenes in both the MVA and methylerythritol phosphate (MEP) pathways (Fig. 4). Transcriptional analysis revealed that most of the terpene biosynthesis genes were more highly expressed in FBs, such as *HMGR*, *HDS*, and *TPS* genes. More importantly, some *TPS* genes regulating synthesis of germacrene (sesquiterpene), geraniol (monoterpene), and alpha-terpineol (monoterpene) were also expressed more highly in FBs than in FFs. These products contribute significantly to floral fragrance. However, three genes encoding TPSs (*JS6G23500*, *JS4G13880*, and *JS4G16830*) responsible for α-farnesene and linalool synthesis were highly expressed in FFs, and metabonomic analysis further revealed that α-farnesene and linalool contents were higher in FFs (Supplementary Tables S10, S11, S12). In addition, several other sesquiterpenes (such as isoledene and cis-caryophyllene) and diterpenes (muurolene) were also detected, all of which had higher levels in FFs (Fig. 4).

**Fig. 4.**
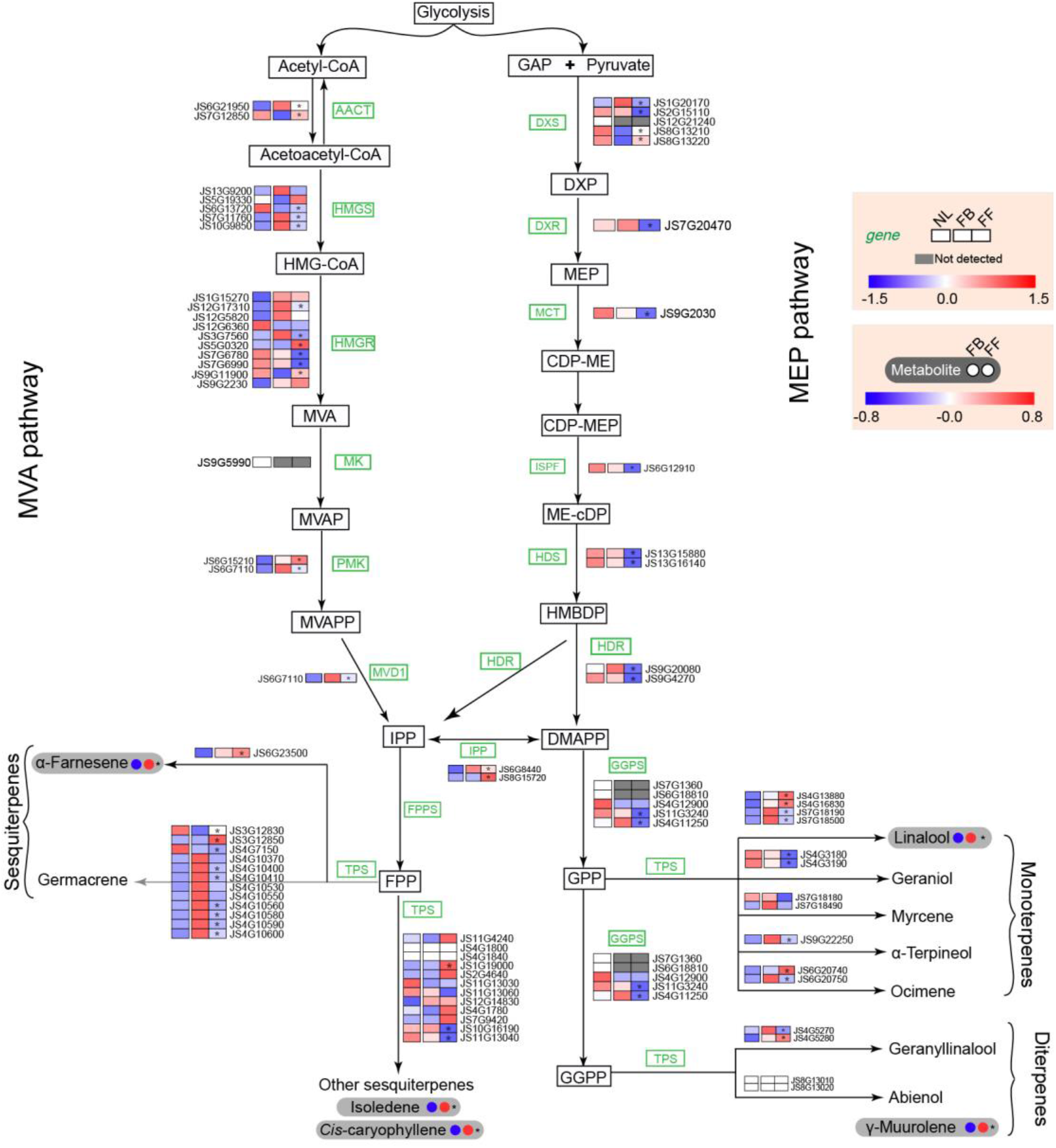
Terpene synthesis pathways in *J. sambac* leaves and flowers based on transcriptomic and metabolomic analyses. Heatmap columns indicate expression levels of genes involved in terpene synthesis pathways. Circles to the right of metabolites highlighted in grey indicate the different metabolite contents of flower buds (left) and full-bloom flowers (right). Asterisks indicate marked differences between the flower buds and full-bloom flowers. NL, normal leaves; FB, flower buds; FF, full-bloom flowers.

### Phenylpropanoid/benzenoid biosynthesis in *J. sambac*

Phenylpropanoids and benzenoids represent the second largest class of flower VOCs^20^ and are exclusively derived from the aromatic amino acid phenylalanine (Phe) (Fig. 5a). Our metabolomic and transcriptomic analyses identified many genes and metabolites involved in the phenylpropanoid/benzenoid pathways. The expression levels of the gene encoding phenylalanine ammonia-lyase (PAL), the first committed enzyme in phenylpropanoid/benzenoid pathways, was higher in FFs than in FBs. Moreover, the expression levels of other genes, including *AAAT*, *EGS*, *IGS*, and *SAMT*, were also higher in FFs, while those of some genes in phenylpropanoid/benzenoid pathways (such as *BPBT*) were lower in FFs (Fig. 5a). The production of phenylpropanoid/benzenoid compounds in plants is related to the SABATH and BAHD acyltransferase super-families. In our analyses, 11 SABATH homologs were identified, belonging to the IAMT (3), SAMT (2), JMT (1), SAMT/BSMT (1), and FAMT-like (4) subfamilies (Fig. 5b). Transcriptomic analysis revealed that expression of FAMT-like genes was higher in FBs than in FFs, while *JMT* and *SAMT* genes were more highly expressed in FFs, and *IAMT* genes were expressed at low levels at both stages (Fig. 5c). In addition, COMT and ICMT, belonging to the SAM-binding methyltransferase superfamily, are involved in aromatic compound metabolism. Our analysis revealed that expression of *COMT* genes was higher in FFs, while that of *ICMT* genes was higher in FBs (Fig. 5c). BAHD acyltransferases are responsible for the synthesis of a myriad flavors and fragrances in plants. In our analysis, expression of most of the genes in the BAHD family was higher in FBs (Fig. 5d). However, expression of most genes in the phenylpropanoid/benzenoid pathways was low in leaves (Fig. 5a, c, d), indicating more active phenylpropanoid/benzenoid biosynthesis in flowers than in leaves. In addition, our metabolomic analysis revealed that a majority of the detected metabolites, including PhEth, PhA, Eug, benzyl benzoate (BB), benzyl alcohol (BAlc), benzyl acetate (BAC), BAld, MB, and methyl salicylate (MeSA), accumulated markedly in FFs, whereas salicylic acid (SA) was higher in FBs (Fig. 5a).

**Fig. 5.**
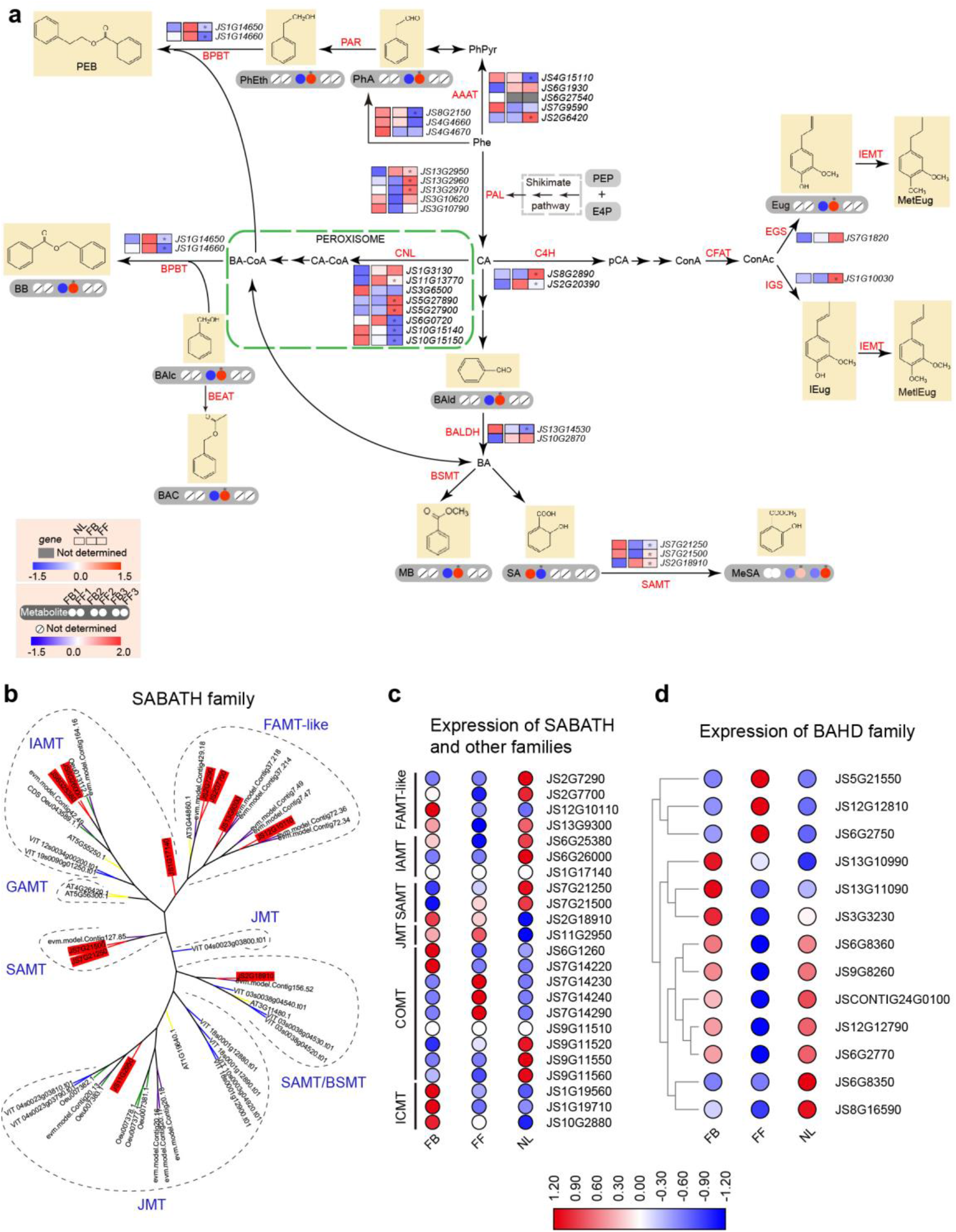
Phenylpropanoid/benzenoid biosynthesis in *J. sambac*. **a** Phenylpropanoid/benzenoid biosynthesis pathways in *J. sambac* leaves and flowers based on transcriptomic and metabolomic analyses. **b** Phylogenetic tree of the SABATH family in plants. The 11 SABATH homologs identified in *J. sambac* are in red. **c** Transcriptomic analysis of SABATH family genes and other families in *J. sambac* leaves and flowers. **d** Transcriptomic analysis of BAHD family genes in *J. sambac* leaves and flowers. NL, normal leaves; FB, flower buds; FF, full-bloom flowers.

### Jasmonate biosynthesis genes

Based on comparisons with the genomes of *A. thaliana*, tomato, *Antirrhinum majus*, and *O. europaea*, no expansion of the jasmonic acid (JA) biosynthesis genes was found in the *J. sambac* genome. However, the number of genes was significantly higher compared to *A. thaliana* (Fig. 6a). Several key genes in the regulation of JA biosynthesis, including *OPR*, *OPCL*, *AOS*, and *KAT,* were present at a ratio of 2:1 relative to their presence in the tomato genome, indicating duplication of these genes in *J. sambac* resulting from the Oleaceae-specific WGD event (Fig. 6b). Transcriptomic analysis revealed that the expression levels of JA biosynthesis genes in FBs and FFs were quite different. The expression levels of genes in the JA biosynthesis pathway, especially *AOS*, *AOC*, *MFP*, and *KAT*, were much higher in FBs than in FFs, while those of β-oxidation genes such as *OPR*, *OPCL*, and *ACX* were higher in FFs (Fig. 6c, e). In the signaling pathway, the expression levels of *JS11G18210* and *JS11G18220* were higher in FBs than in FFs, while those of *JS7G16870*, *JS10G13760*, *JS10G200*, *JS5G30430*, and *JS4G4500* were higher in FFs (Fig. 6c). In addition, metabolomic and liquid chromatography-mass spectrometry (LC-MS) analyses both showed that jasmonates were enriched in FFs and FBs. The contents of JA, methyl jasmonate (MeJA), and jasmonic acid-isoleucine (JA-Ile), as well as those of their precursor (α-linolenic acid) and intermediate metabolite (12-oxo-phytodienoic acid, OPDA), were higher in FFs than in FBs (Fig. 6c, d, Supplementary Table S13). Moreover, LC-MS analysis also revealed that JA and JA-Ile contents were significantly higher in leaves at 20 min to 2 h after wounding (Fig. 6d). Interestingly, some genes with high expression in flowers had low expression in leaves, but these genes were significantly highly expressed in wounded leaves (Fig. 6c, e). Specifically, the mean expression levels of JA synthesis-related genes (*AOS*, *ACX1*, *KAT*) were higher in wounded leaves than in normal leaves, especially at 1–2 h after wounding, and JA signal-transduction-related genes (*JAZ*, *MYC2*) were activated at 20 min to 1 h after wounding (Fig. 6e). Quantitative reverse-transcription polymerase chain reaction (qRT-PCR) analysis also demonstrated that the expression of JA synthesis-related genes (*AOC*, *AOS*, *OPR*) and JA signal-transduction-related genes (*JAZ*, *MYC2*) significantly increased in leaves at 1–2 h after wounding, and then decreased (Fig. 6f). Of note, the expression levels of *JS5G30430* (one of the *JAZs*) were low in both normal leaves and wounded leaves. However, it was highly expressed in FFs, indicating that *JS5G30430* mainly responds to endogenous signals during flower development rather than participating in the JA-mediated injury response.

**Fig. 6.**
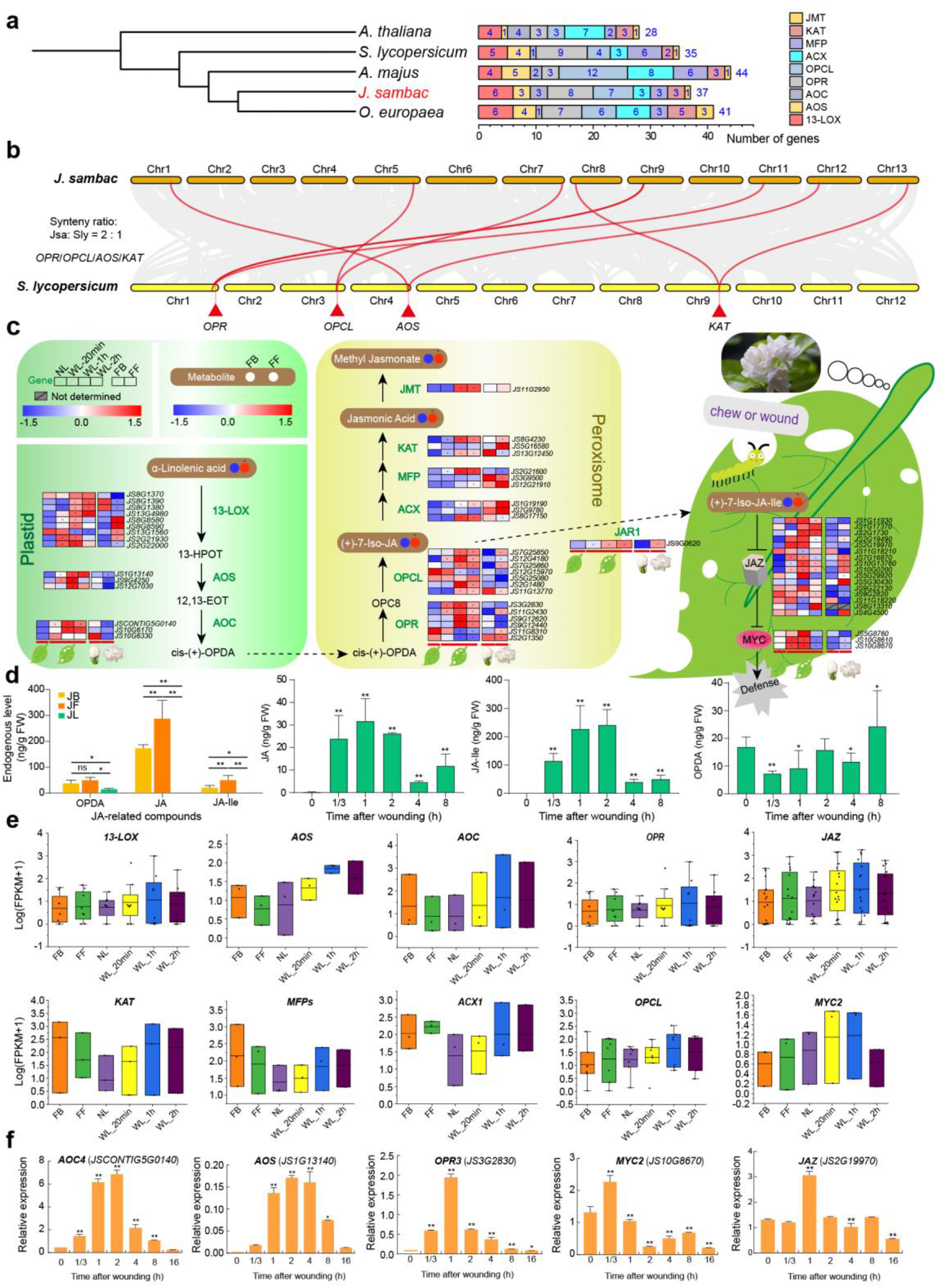
Jasmonate biosynthesis in *J. sambac*. **a** Comparison of jasmonate biosynthesis genes in the genomes of *J. sambac*, *A. thaliana*, *S. lycopersicum*, *A. majus*, and *O. europaea*. **b** Synteny analysis in genomes of *J. sambac* and *S. lycopersicum.* **c** Jasmonate synthesis pathways in *J. sambac* normal leaves, wounded leaves, and flowers based on transcriptomic and metabolomic analyses. **d** Contents of endogenous JA-related compounds in flowers and wounded leaves. **e** Transcriptomic analysis of genes involved in jasmonate biosynthesis and JA signal transduction in normal leaves, wounded leaves, and flowers. **f** qRT-PCR analysis of genes involved in jasmonate biosynthesis and JA signal transduction in leaves at different times after wounding. NL, normal leaves; WL, wounded leaves (followed by time after wounding); FB, flower buds; FF, full-bloom flowers.

## Discussion

*J. sambac* is the world-famous flower that is widely used in ornamental horticulture, the perfume industry, scented tea, food, and pharmaceutical applications^2,21,22^. Despite its extensive use, the genome of *J. sambac* has not yet been sequenced. In this study, we used the cultivar ‘double petal’ (the most widely-cultivated *J. sambac*) as material for genome sequencing (Fig. 7a, b). We sequenced and assembled a high-quality chromosome-level genome of *J. sambac* by combining PacBio and Illumina with Hi-C sequencing. Our assembled genome is approximately 550.12 Mb with a scaffold N50 size of 40.10 Mb; 97.36% of the genome is anchored onto 13 pseudochromosomes. Furthermore, BUSCO evaluation revealed that the genome covers 91.7% of the complete single-copy orthologs of plant-specific sequences. These results indicate that our genome assembly is precise, complete, and of high quality. In addition, 30,129 genes were annotated by the combination of *de novo*, homology-based, and RNA sequencing (RNA-seq) data, 93.2% of which had predicted functions, indicating a high annotation quality. This high-quality genome sequence of *J. sambac* provides a fundamental genetic resource for functional genomic research and understanding fragrance biosynthesis mechanisms in *J. sambac*.

**Fig. 7.**
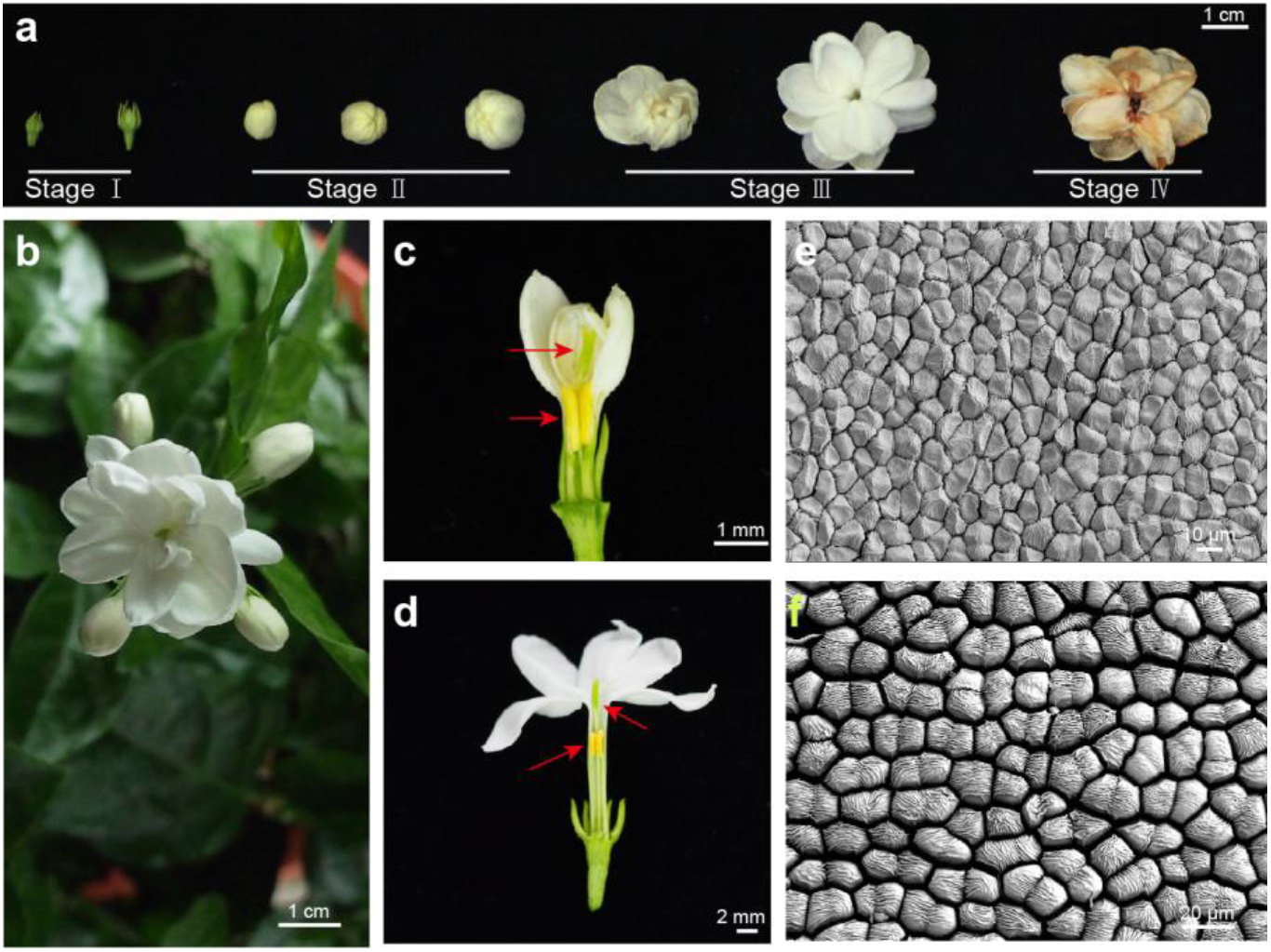
Morphology of *J. sambac* flowers. **a** Different stages of *J. sambac* flowers. **b** The morphology of fully blooming flowers. **c, d** Anatomy of *J. sambac* flower bud and full-bloom flower. Red arrows indicate the pistil and stamens. **e, f** Morphology of the petal cells of a flower bud (e) and full-bloom flower (f) under scanning electron microscopy.

In the Oleaceae family, the genomes of five species have been sequenced, including *F. excelsior*^23^, *O. europaea*^24^, *O. oleaster*^25^, *O. fragrans*^26^, and *Forsythia suspensa*^27^. Among them, only *O. fragrans* flowers are scented with a sweet aroma, with VOCs including linalool, dihydrojasmone lactone, 1-cyclohexene-1-propanol, and β-ocimene^28^. By contrast, the aroma type of *J. sambac* flowers is significantly different. During the flowering period, as their petals fully expand, the corolla tube in FFs becomes markedly elongated compared to that in FBs (Fig. 7c, d), indicating that this developmental stage is for flower fragrance release. We identified the major volatile fragrances in *J. sambac* flowers as terpenoids (linalool, γ-muurolene, isoledene, farnesene), phenylpropanoids/benzenoids (phenylacetaldehyde, 2-phenylethanol, BB, BAlc, BAC, MeSA), and fatty acids (α-linolenic acid, JA, MeJA, JA-Ile) (Supplementary Tables S11, S13). These VOCs appear to make up the unique scent of *J. sambac* flowers. Generally, plant VOC biosynthesis is regulated by many genes and gene families^20,29^. For example, the terpenoids are biosynthesized via TPS-dependent pathways^30^. In the genome of *J. sambac*, we identified many TPS genes present as gene clusters through recent tandem duplications, resulting in significantly amplified TPS genes in the genome. Therefore, the terpenoid fragrance enriched in *J. sambac* flowers is likely contributed by these TPS gene clusters. Moreover, the terpenoid fragrance that evaporates into the air differs significantly between FBs and FFs. This can be explained by the differential expression of TPS genes observed at the two stages. In addition, FF petal cells are plump, with larger intercellular spaces, compared to those in FBs (Fig. 7e, f), implying that the separated petal cells increase the emission of fragrant compounds in FFs. These morphological, transcriptional, and metabolomics analyses collectively imply that fragrance release from *J. sambac* flowers is a dynamic stage-dependent process. Notably, negative selection occurs in many TPS gene pairs in the *J. sambac* genome. Since *J. sambac* has already been cultivated for over 1000 years^31^, this type of negative selection has largely resulted from long-term selection through artificial cultivation, especially by vegetative propagation in *J. sambac*.

Volatile phenylpropanoids and benzenoids are major volatile aromas present in plants^10^. They originate from the aromatic amino acid Phe. Several other antioxidant metabolites are also synthesized from Phe, including flavonoids and anthocyanin pigments^32,33^. Phe is deaminated to cinnamic acid (CA) by PAL. CA is further converted into diverse volatile compounds via three main synthetic routes of enzymatic and acid-catalyzed transformations: BB, catalyzed by cinnamate-coenzyme A ligase (CNL); Eug, catalyzed by EGS; and MB, SA, and MeSA, catalyzed by benzaldehyde dehydrogenase (BALDH). Phe can also be converted to PEB via PhA and PhEth, catalyzed by PAR and BPBT, respectively^20^. In our analysis, these phenylpropanoid/benzenoid volatile compounds, including BB, Eug, MB, MeSA, PhA, and PhEth (Supplementary Table S14), accumulated more in FFs than in FBs, whereas SA was detected at higher levels in FBs (Fig. 5a). Furthermore, many other metabolites in the phenylpropanoid/benzenoid pathways were also detected in our analysis, such as phenylpyruvic acid, ferulic acid, 2,3-dihydroxybenzoic acid, and benzoic acid. These volatile compounds also contribute to the specific aromas of *J. sambac* flowers. Expression analysis of the phenylpropanoid/benzenoid pathway genes by RNA-seq revealed that *PAL*, *AAAT*, *EGS*, *IGS*, and *SAMT* were highly expressed in FFs, whereas *BPBT* and *CNL* were more highly expressed in FBs. Apparently, the regulation of gene expression at varying stages coordinates VOC dynamics during the timeline of flower blooming. Notably, we identified many flavonoids (38 of 174) that were enriched in *J. sambac* flowers. As flavonoids are important secondary metabolites with antioxidant properties, and jasmine tea is a common beverage, the jasmine flowers in tea may be beneficial to human health in addition to providing aroma.

Another important type of fragrant VOCs in *J. sambac* flowers is fatty acids and their derivatives. Among them, jasmonate and its related compounds, including JA, MeJA, JA-Ile, and jasmone, are fragrant components of the essential oils of jasmine (*Jasminum*) flowers^2,13^. In our metabolomics analyses, these jasmonate-related compounds were enriched in *J. sambac* flowers, indicating their important roles in the formation of the characteristic aromatic odor of the flowers. In addition, JA, MeJA, and JA-Ile play important roles in plant defense against biotic and abiotic stresses^18^. Our analysis revealed that the JA contents and related genes in *J. sambac* leaves responded to mechanical injury. For example, several genes in the JA biosynthesis pathway were highly expressed in wounded leaves and the JA signal-transduction-related genes (*JAZ*, *MYC2*) were also activated after wounding. These results were further confirmed by qRT-PCR. Compared to *Arabidopsis* and tobacco^34,35^, the response times of genes involved in JA signaling and biosynthesis pathways to wounding are similar in leaves, implying a similar responsive pattern in *J. sambac.* However, in the JA signaling pathway, expression of some *JAZ* genes, such as *JS11G18210* and *JS11G18220*, was significantly higher in FBs, while that of other *JAZ* genes (*JS7G16870*, *JS10G13760*, *JS10G200*, *JS5G30430*, and *JS4G4500*) was higher in FFs, implying a significant difference in the function of the *JAZs* involved in the flower development of *J. sambac*. Moreover, *JAZs* and *MYCs* were significantly activated with the high JA content in FFs, implying that flowering is the important period for biosynthesis of jasmonates and JA-related floral aroma substances. These findings demonstrate that the regulation of the JA signaling pathway during *J. sambac* flower development is related to the robust secondary metabolism. In particular, MeJA has been reported to induce expression of terpenoid-and phenylpropanoid/benzenoid-related genes and promote the synthesis of related metabolites^32,36^. Therefore, these jasmonates in blooming *J. sambac* flowers may also affect the synthesis and release of other floral aromatic components. As an important aroma itself, MeJA together with other aromatic components may orchestrate the unique sweet fragrance of *J. sambac* flowers. However, other than attracting pollinators, the further biological significance of jasmonates in *J. sambac* flowers requires additional research.

In summary, we here present a chromosome-level genome of *J. sambac* and identify the main volatile aromas in *J. sambac* flower buds and blooming flowers. Our multi-omics analyses reveal the mechanisms of jasmine volatile aroma production (Fig. 8). This high-quality, annotated genome sequence of *J. sambac* together with the transcriptomic and metabolomic datasets in this study provide a fundamental genetic resource for studying functional genomics and fragrance biosynthesis in *J. sambac*, which will be invaluable for industrial exploitation of jasmine flowers in the future.

**Fig. 8.**
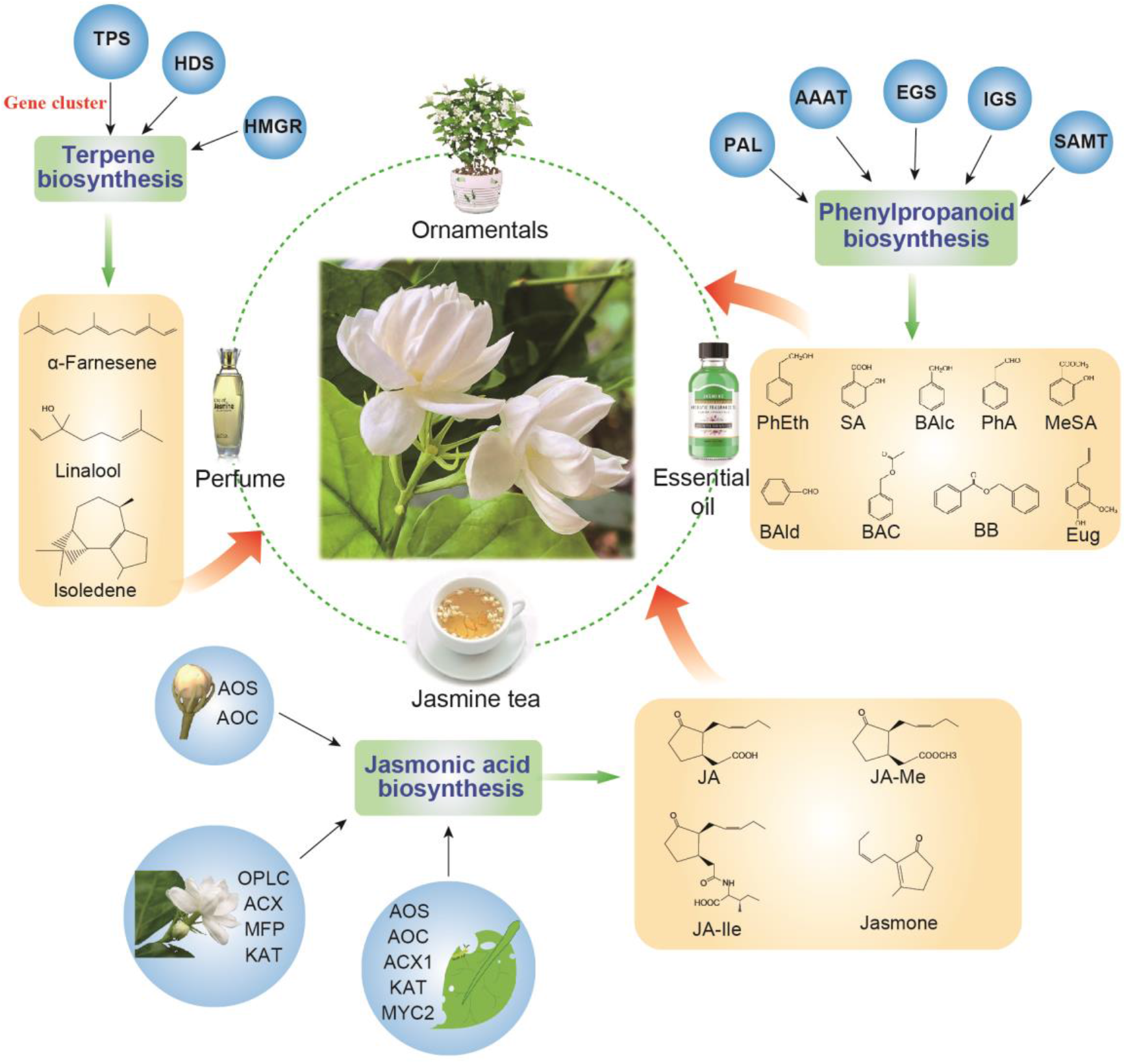
Schematic diagram summarizing the production of aromatic compounds in *J. sambac* flowers and their contributions to commercial applications of *J. sambac.*

## Materials and Methods

### Plant materials for genome sequencing

The *J. sambac* ‘double petal’ cultivar, the major cultivar in China, was selected as the model plant species for studying jasmine flowers (Fig. 7). All the materials were sampled from individual potted plant clones with the same genetic background in the greenhouse of Yangzhou University, Yangzhou, China (32.39° N, 119.42° E). The newly expanded leaves from the sequenced plant were disinfected with 70% ethyl alcohol and rinsed with distilled water. They were then harvested and immediately frozen in liquid nitrogen and stored at −80°C prior to DNA extraction.

### Estimation of the genome size

To estimate the *J. sambac* genome size, k-mer analysis with Illumina sequencing short reads was performed. A k-mer refers to an oligonucleotide of k bp in length. The k-mer frequencies derived from the sequencing reads follow a Poisson distribution in a given dataset (Supplementary Fig. S1). Given a certain k-mer, genome size can be simply inferred from the total number of k-mers (referred to as K_num) divided by the k-mer depth: genome size = K_num / k-mer_depth. When the k-mer size was set to 17, the 89.05 Gb (sequencing depth of 162×) of sequencing reads from short-insert size libraries generated a total of 64,963,845,770 k-mers, and the k-mer depth was ~112×. From these statistics, we estimated that the genome size of *J. sambac* was ~580.03 Mb, and after removing the wrong k-mers, the revised genome size was ~573.02 Mb.

### Library preparation and sequencing

Genomic DNA was extracted by the CTAB method and at least 10 μg of sheared DNA was obtained. SMRT bell template preparation, including DNA concentration, damage repair, end repair, ligation of hairpin adapters, and template purification, was conducted using AMPure PB Magnetic Beads (PacBio, Menlo Park, CA, USA). We conducted 20-kb single-molecule real-time DNA sequencing using PacBio to sequence a DNA library on the PacBio Sequel platform and 63.90 Gb of raw sequencing reads were obtained. Reads were trimmed for adaptor removal and quality enhancement, yielding 63.82 Gb of PacBio data (read quality ≥ 0.80, mean read length ≥ 14 kb) representing 116× genome coverage.

### De novo *genome assembly*

Before *de novo* assembly, low-quality PacBio subreads with a read length < 500 bp or a quality score < 0.8 were filtered out. The remaining clean PacBio subreads were error-corrected and assembled into contigs using FALCON software. The assembled scaffolds were polished with Quiver (http://pbsmrtpipe.readthedocs.io/en/master/getting_started.html) in two rounds. Finally, the polished sequences were further corrected with reference to the Illumina reads using Pilon (https://github.com/broadinstitute/pilon/wiki) in two rounds.

### Pseudochromosome validation using Hi-C

To avoid artificial bias, the following reads were removed: reads with ≥ 10% unidentified nucleotides (Ns); reads with > 10 bp aligned to the adapter, allowing ≤ 10% mismatches; reads with > 50% bases having a Phred quality < 5. The filtered Hi-C reads were aligned to the initial pseudochromosome genome using BWA (version 0.7.8) with default parameters. Reads were excluded from subsequent analysis if they did not align within 500 bp of a restriction site. Only paired-end reads that uniquely mapped with valid ditags were used to validate the pseudochromosome sequences. Juicebox (https://github.com/aidenlab/Juicebox) was used to manually order the scaffolds in each group to obtain the final pseudochromosome assembly. Contact maps were plotted using HiCPlotter. The high collinearity between the genetic map-based chromosome anchoring and Hi-C-based contact map information corroborated the overall assembly quality.

### *Assessment of* J. sambac *genome quality*

The completeness of the *J. sambac* assembly was evaluated by two methods. First, the 1440 conserved protein models in the BUSCO “embryophyta_odb9” dataset (https://busco.ezlab.org/frame_wget.html) were queried against the *J. sambac* genome using the BUSCO (Version 2) program with default settings (Supplementary Table S3); we obtained a genome completeness value of 91.7%.

### Repeat annotation

Repeat elements in the *J. sambac* genome were annotated using a combined strategy. Alignment searches were undertaken against the Repbase database (http://www.girinst.org/repbase), then RepeatProteinMask searches (http://www.repeatmasker.org/) were used for prediction of homologs^37^. For *de novo* annotation of repeat elements, LTR_FINDER (http://tlife.fudan.edu.cn/tlife/ltr_finder/) Piler (http://www.drive5.com/piler/), RepeatScout (http://www.repeatmasker.org/), and RepeatModeler (http://www.repeatmasker.org/RepeatModeler/) were used to construct a *de novo* library, then annotation was carried out with RepeatMasker.

### RNA-seq-based prediction for gene annotation

To aid gene annotation and perform the transcriptome analysis, RNA was extracted from six tissues (roots, shoots, adult leaves, wounded leaves, buds, and full-bloom flowers) of *J. sambac*. All fresh tissues were first frozen in liquid nitrogen and stored at −80°C before processing. Total RNA of each sample was extracted using TRIzol Reagent (Invitrogen, Carlsbad, CA, USA) according to the manufacturer’s instructions and mixed together. RNA-seq libraries were prepared using the Illumina standard mRNA-seq library preparation kit and sequenced on the Illumina HiSeq 4000 platform using a paired-end sequencing strategy. A full-length isoform sequencing (ISO-seq) library was also constructed with an insert size of 0–5 kb using the same samples, then sequenced on the PacBio SMRT Sequel platform at Novogene (Tianjin, China). The ISO-seq reads were extracted using the SMRTlink (https://www.pacb.com/support/software-downloads/) software to obtain the polished consensus sequences; these data were further processed by the CD-hit software to remove redundancies.

### Annotation of protein-coding genes

Protein-coding genes were annotated using a comprehensive strategy integrating results obtained from homology-based prediction, *de novo* prediction, and RNA-seq-based prediction methods. Annotated protein sequences from *F. excelsior*, *O. fragrans*, *O. europaea*, and *O. europaea* var. *sylvestris* (Oleaceae family) were aligned to the *J. sambac* genome assembly using WU-Blast with an E-value cutoff of 1e^−5^ and the hits were conjoined using the Solar software. GeneWise was used to predict the exact gene structure of the corresponding genomic regions for each WU-Blast hit. The gene structure created by GeneWise was denoted as the homology-based prediction gene set (Homo-set). Gene models created by PASA were denoted as the PASA ISO-seq set (PASA-ISO-set) and were used as the training data for the *de novo* gene prediction programs. Five *de novo* gene-prediction programs (Augustus, GENSCAN, GeneID, GlimmerHMM, and SNAP) were used to predict coding regions in the repeat-masked genome. Illumina RNA-seq data were mapped to the assembly using TopHat, then Cufflinks was used to assemble the transcripts into gene models (Cufflinks-set). In addition, RNA-seq data were assembled by Trinity, creating several pseudo-expressed sequence tags (ESTs). These pseudo-ESTs were also mapped to the SCHZ assembly by LASTZ, and gene models were predicted using PASA. PacBio ISO-seq sequences were mapped directly to the *J. sambac* genome assembly by BLAT and assembled by PASA. This gene set was denoted as the PASA Trinity set (PASA-T-set). Gene model evidence data from the Homo-set, PASA-ISO-set, Cufflinks-set, PASA-T-set, and *de novo* programs were combined by EvidenceModeler into a non-redundant set of gene annotations. Weights for each type of evidence were set as follows: PASA-ISO-set > Homo-set > PASA-T-set > Cufflinks-set > Augustus > GeneID = SNAP = GlimmerHMM = GENSCAN. Gene models with low confidence scores were filtered out by the following criteria: (1) coding region lengths of 150 bp, (2) supported only by *de novo* methods and with FPKM<1.

All protein-coding genes were aligned to two integrated protein sequence databases: SwissProt and NR. Protein domains were annotated by searching against InterPro database (Version 32.0) using InterProScan and against the Pfam database (Version 27.0) using HMMER. The GO terms for each gene were obtained from the corresponding InterPro or Pfam entry. The pathways in which the genes might be involved were assigned by BLAST searches against the KEGG database, with an E-value cutoff of 1e^−5^. Functional annotation results from the three strategies above were finally merged. The annotation results can be found in Supplementary Table S5. In total, 30,129 genes were predicted to be functional, accounting for 93.2% of all genes in the *J. sambac* genome (Supplementary Table S6).

### Annotation of non-coding RNAs

Non-coding RNAs were annotated using tRNAscan-SE (http://lowelab.ucsc.edu/tRNAscan-SE/) (for tRNA) or INFERNAL (http://infernal.janelia.org/) (for miRNA and small nuclear RNA). Since rRNA sequences are highly conserved among plants, rRNA from *A. thaliana* was screened by Blast searches (Supplementary Table S7).

### Protein ortholog analysis

Orthologous relationships between genes of *J. sambac*, *O. fragrans*, *F. excelsior*, *O. europaea*, *O. oleaster*, *Prunus mume*, *Petunia inflata*, *C. sinensis*, *Populus trichocarpa*, *S. lycopersicum*, *Vitis vinifera*, *Medicago truncatula*, *Oryza sativa*, *A. thaliana*, *A. majus*, *Amborella trichopoda*, and *Salvia splendens* were inferred through all-against-all protein sequence similarity searches using OrthoMCL (http://orthomcl.org/orthomcl/); only the longest predicted transcript per locus was retained.

### Protein phylogenetic analysis

For each gene family, an alignment was produced using MUSCLE (http://www.drive5.com/muscle/), ambiguously aligned positions were trimmed using Gblocks (http://molevol.cmima.csic.es/castresana/Gblocks.html), and the tree was inferred using RAxML 7.2.9 (http://sco.h-its.org/exelixis/software.html).

### Estimates of divergence times

Divergence times between species were calculated using the MCMC tree program (http://abacus.gene.ucl.ac.uk/software/paml.html) implemented in Phylogenetic Analysis by Maximum Likelihood (PAML).

### Expansion and contraction of gene families

To identify gene family evolution as a stochastic birth and death process in which a gene family either expands or contracts per gene per million years independently along each branch of the phylogenetic tree, we used the likelihood model originally implemented in the software package Café (http://sourceforge.net/projects/cafehahnlab/). The phylogenetic tree topology and branch lengths were taken into account to infer the significance of changes in gene family size in each branch.

### WGD analysis

We applied 4DTv and *Ks* estimation to detect WGD events. First, respective paralogs of *O. fragrans*, *Glycine max*, *O. europaea*, *V. vinifera*, and *A. thaliana* were identified with OrthoMCL. Then, the protein sequences of these plants were aligned against each other using Blastp (E-value ≤ 1e^−5^) to identify the conserved paralogs of each plant. Finally, the WGD events of each plant were evaluated based on their 4DTv distributions (Fig. 2d).

### Transcriptome analysis of different tissues

The adult leaves were prodded with needles to simulate insect biting; 20 minutes after prodding, the wounded and normal leaves as well as FBs and FFs were collected and stored in liquid nitrogen, then transferred to a freezer at −80°C before RNA extraction. Three biological replicates (each treatment and each tissue) were conducted. RNA-seq libraries were constructed according to the manufacturer’s instructions and sequenced on the Illumina HiSeq 4000 platform at Novogene. The RNA-seq reads of each sample were mapped to the reference genome of *J. sambac* by HISAT2 with the parameter “--dta”; StringTie was further used to calculate the transcript per million (TPM) value for each gene with default parameters.

In addition, the extracted RNAs from wounded leaves, normal leaves, stems, roots, FBs, and FFs were mixed for RNA-seq and full-length transcriptome sequencing to assist with gene annotation.

### Detection of metabolites in FBs and FFs by ultra-performance (UP) LC-MS

The FBs and FFs (Fig. 7a, stages Ⅱ and Ⅲ) were collected and stored in liquid nitrogen, then transferred to a freezer at −80°C. The freeze-dried buds and flowers were crushed using a mixer mill (MM 400, Retsch, Haan, Germany) with a zirconia bead for 1.5 min at 30 Hz. Powder (100 mg) was weighed and extracted overnight at 4°C with 0.6 mL 70% aqueous methanol. Following centrifugation at 10,000 *g* for 10 min, the extracts were absorbed (CNWBOND Carbon-GCB SPE Cartridge, 250 mg, 3 mL; ANPEL, Shanghai, China) and filtrated (SCAA-104, 0.22 μm pore size; ANPEL) before UPLC-MS/MS analysis. Then, the sample extracts were analyzed using a UPLC–electrospray ionization (ESI)-MS/MS system (UPLC: Shim-pack UFLC CBM30A; Shimadzu, Kyoto, Japan; MS: QTRAP 4500; Applied Biosystems, Foster City, CA).

### Detection of volatiles using gas chromatography (GC)-MS

The buds and flowers stored at −80°C were ground to powder in liquid nitrogen. The powder (1 g) was immediately transferred to a 20-mL head-space vial (Agilent, Palo Alto, CA, USA) containing 2 mL NaCl-saturated solution to inhibit any enzyme reaction. The vials were sealed using crimp-top caps with TFE-silicone headspace septa (Agilent). At the time of solid phase microextraction analysis, each vial was incubated at 60°C for 10 min, then a 65-μm divinylbenzene/Carboxen/polydimethylsiloxane fiber (Supelco, Bellefonte, PA, USA) was exposed to the headspace of the sample for 20 min at 60°C. After sampling, desorption of the VOCs from the fiber coating was carried out in the injection port of the GC apparatus (Model 7890B; Agilent) at 250°C for 5 min in splitless mode. The identification and quantification of VOCs was carried out using a 7890B GC and a 7000D mass spectrometer (Agilent) equipped with a 5% phenyl-polymethylsiloxane capillary column (DB-5MS, 30 m × 0.25 mm × 1.0 μm; Agilent).

### Detection of volatiles actively released from flowers on living plants

MonoTrap (DCC 18; Shimadzu) disks were used as absorbents for volatile collection. The overground parts of *J. sambac* plants were covered and fastened to a Teflon gas sampling bag (5 L), with MonoTrap disks hanging on branches next to blooming flowers (Supplementary Fig. S9).

After 6 h of absorption, all MonoTrap disks were collected in sealed bottles (one disk per bottle). The disks were crushed under liquid nitrogen, then carbon disulfide was used to elute and collect absorbed volatiles. Then, the Exactive GC Orbitrap GC-MS system (Thermo Fisher Scientific, Waltham, MA, USA) coupled with a Tace1310 GC was used for metabolite analysis.

Extracts were liquid injected and separated by a DB-5 column using the following GC program: start at 40°C, hold for 5 min, then increase temperature to 280°C at a rate of 5°C/min. The scan range of 33–550 m/z was acquired with data dependent MS/MS acquisition with 60,000 resolution under full scan mode. The source parameters were as follows: ion source temperature: 280°C; MS transfer line: 250°C.

MS/MS data were analyzed using TraceFinder analysis software (Thermo Fisher Scientific). Data processing parameter settings were as follows: minimum peak width = 10 s, maximum peak width = 60 s, mzwid = 0.015, minfrac = 0.5, bw = 5, and signal/noise threshold = 5.

Metabolites were identified and characterized based on high-resolution MS-associated methods using TraceFinder. First, the candidate chemical formulas of metabolites were identified using the accurate high-resolution m/z values and MS/MS fragment patterns, with a mass accuracy of 3 ppm based on NIST MS Search. Then, the MS/MS spectra were analyzed by manually comparing both fragment patterns and isotope ratios to identify the metabolites. Peak detection, retention time correction, and chromatogram alignment were performed. The results contained a peak list with metabolite names, retention times, m/z values, and the mean ion abundance with standard deviation.

### Plant hormone extraction

Flower and leaf samples were collected and immediately stored in liquid nitrogen. Then, metabolites of samples were extracted using a modified Wolfender method. First, 100 mg of flower powder obtained by crushing under liquid nitrogen was weighed and transferred to a 2-mL centrifuge tube with 10 μL internal standards (10 μg/mL d5-JA). Second, 1.5 mL extraction buffer (isopropanol:formic acid = 99.5:0.5, v/v) was added followed by vortexing to resuspend samples. After 15 min centrifugation at 14,000 *g*, the supernatants were dried in a Labconco CentriVap vacuum centrifugal concentrator and resuspended with 1 mL methanol solvent (85:15, v/v). Then, a C18 SPE tube (Sep-pak C18 SPE Cartridge, 100 mg, 1 mL; Waters Technology, Shanghai, China) was used for sample purification, and a total of 1.5 mL eluent was collected for each sample. Finally, the eluents were dried in a Labconco CentriVap vacuum centrifugal concentrator and resuspended with 100 μL methanol solvent (60:40, v/v).

### LC-MS for plant hormones

Positive/negative ionization mode data were acquired using an Acquity UPLC I-Class (Waters Technology) coupled to a 4500 QTRAP triple quadrupole mass spectrometer (AB SCIEX, Ontario, CA) equipped with a 50 × 2.1 mm, 1.7 μm Acquity UPLC BEH C18 column (Waters Technology); 10-μL samples were loaded each time, and then eluted at a flow rate of 200 μL/min with initial conditions of 50% mobile phase A (0.1% formic acid in acetonitrile) and 50% mobile phase B (0.1% formic acid in water) followed by a 10-min linear gradient to 100% mobile phase A. The auto-sampler was set at 10°C.

Mass spectrometry was operated separately in positive/negative ESI mode. The [M+H] or [M−H] of the analyte was selected as the precursor ion; precursor ion/product ion details for quantitation under multiple reaction monitoring mode are shown in Supplementary Table S15. The temperature of the ESI ion source was set to 500°C. Curtain gas flow was set to 25 psi, collisionally activated dissociation gas was set to medium, and the ionspray voltage was (+)5500 V for positive ionization mode and (−)4500 for negative ionization mode with ion gases 1 and 2 set to 50 psi. Data acquisition and processing were performed using AB SCIEX Analyst version 1.6.3 (Applied Biosystems)

## Data availability

The genome assemblies, gene annotations, and Illumina re-sequencing short reads have been deposited in the Genome Sequence Archive (GSA) and Genome Warehouse database in the BIG Data Center (https://bigd.big.ac.cn/gsa/) under BioProject Accession number GSA:PRJCA003967.

## Acknowledgements

We thank Novogene for genome sequencing and assembly. We thank Dr. Feng Cheng for his comments on our manuscript. This work was self-funded.

## Author notes

These author contributed equally: Gang Chen, Salma Mostafa, Zhaogeng Lu, Ran Du.

## Contributions

B.J., J.Y, and G.C. conceived and designed the study; G.C., S.M., Z.L., R.D., J.C., Y.W., Q.L., J.L., X.M., B.C., L.W., and Z.J. performed the experiments; G.C., S.M., Z.L., R.D., J.C., Y.W., Q.L., J.L., X.M., B.C., L.W., Z.J., X.Y., and Y.Z. analysed the data; B.J., J.Y, and G.C. wrote the manuscript; All authors read and approved the final draft.

## Corresponding authors

Correspondence to Biao Jin or Jianbin Yan.

## Ethics declarations

Not applicable.

## Additional information

### Supplementary Figures

Supplementary Fig. S1. Evaluation of the genome size of *J. sambac* by 17-mer analyses.

Supplementary Fig. S2. Hi-C interaction heatmap of *J. sambac* reference genome showing interactions between the 13 chromosomes.

Supplementary Fig. S3. Divergence distribution of transposable elements in the genome of *J. sambac*.

Supplementary Fig. S4. Different elements of annotated genes in *J. sambac* genome.

Supplementary Fig. S5. Summary of protein-coding genes in *J. sambac* genome predicted from *de novo*, homology-based, and RNA-seq approaches.

Supplementary Fig. S6. Functional annotation of genes in *J. sambac* genome.

Supplementary Fig. S7. KEGG enrichment results of the *J. sambac* specific gene families.

Supplementary Fig. S8. KEGG enrichment of different expressed genes in flower buds and full-blown flowers (a) and synteny analysis in genomes of *J. sambac* and *S. lycopersicum* (b).

Supplementary Fig. S9. Volatiles collection with MonoTrap® DCC 18.

### Supplementary Tables

Supplementary Table S1 Genome assembly using both PacBio reads and Hi-C data.

Supplementary Table S2 Distribution of chromosomes in the assembled genome of *J. sambac*.

Supplementary Table S3 Quality assessment of the assembled genome of *J. sambac* using BUSCOs.

Supplementary Table S4 Summary statistics of the annotated transposable elements (TEs) in the *J. sambac* genome.

Supplementary Table S5 Summary statistic of annotated genes in *J. sambac* genome. Genes were annotated by the combination of *de novo*, homology-based, and RNA-seq data.

Supplementary Table S6 Summary statistics of the functional genes of *J. sambac*.

Supplementary Table S7 Annotated non-coding RNA in *J. sambac* genome.

Supplementary Table S8 Functional annotation of the genes in *J. sambac*.

Supplementary Table S9 TPS genes containing at least one conserved domain in the genome of *J. sambac*.

Supplementary Table S10 Volatile metabolites in flower buds and full-bloom flowers in *J. sambac*.

Supplementary Table S11 GC-MS results in flower buds and full-bloom flowers in *J. sambac*.

Supplementary Table S12 Differential abundant terpene metabolites identified between the flower buds and full-bloom flowers in *J. sambac*.

Supplementary Table S13 Widely-targeted metabolomics in flower buds and full-bloom flowers in

*J. sambac*.

Supplementary Table S14 Differential aboundant phenylpropanoid/benzenoid metabolites identified between the flower buds and full-bloom flowers in *J. sambac*.

Supplementary Table S15 The collision energies for different MRM pairs.

